# Separation of telomere protection from length regulation by two different point mutations at amino acid 492 of RTEL1

**DOI:** 10.1101/2024.02.26.582005

**Authors:** Riham Smoom, Catherine Lee May, Dan Lichtental, Emmanuel Skordalakes, Klaus H. Kaestner, Yehuda Tzfati

**Affiliations:** Department of Genetics, The Silberman Institute of Life Sciences, Safra Campus, The Hebrew University of Jerusalem, Jerusalem, 91904, Israel; Department of Genetics and Institute for Diabetes, Obesity and Metabolism, Perelman School of Medicine, University of Pennsylvania, Philadelphia, PA 19104, USA; Department of Pharmacology and Toxicology, Massey Cancer Center, Virginia Commonwealth University, 401 College St, Richmond, VA 23298, USA

## Abstract

RTEL1 is an essential DNA helicase that plays multiple roles in genome stability and telomere length regulation. A variant of RTEL1 with a lysine at position 492 is associated with short telomeres in *Mus spretus*, while a conserved methionine at this position is found in *M. musculus*, which has ultra-long telomeres. In humans, a missense mutation at this position (*Rtel1*^M492I^) causes a fatal telomere biology disease termed Hoyeraal-Hreidarsson syndrome (HHS). Introducing the *Rtel1*^M492K^ mutation into *M. musculus* shortened the telomeres of the resulting strain, termed ‘Telomouse’, to the length of human telomeres. Here, we report on a mouse strain carrying the *Rtel1*^M492I^ mutation, termed ‘HHS mouse’. The HHS mouse telomeres are not as short as those of Telomice but nevertheless they display higher levels of telomeric DNA damage, fragility and recombination, associated with anaphase bridges and micronuclei. These observations indicate that the two mutations separate critical functions of RTEL1: M492K mainly reduces the telomere length setpoint, while M492I predominantly disrupts telomere protection. The two mouse models enable dissecting the mechanistic roles of RTEL1 and the different contributions of short telomeres and DNA damage to telomere biology diseases, genomic instability, cancer, and aging.

## INTRODUCTION

Telomeres are repetitive sequences bound by specialized proteins that protect chromosome ends from deleterious activation of the canonical DNA damage response (DDR) (reviewed in (1)). In all vertebrates, telomeres consist of 5’-TTAGGG-3’ DNA repeats, and end with an essential single-stranded 3’-overhang. This overhang can be elongated by the enzyme telomerase to compensate for sequence loss due to the inability of the canonical replication machinary to fully replicate the chromosome end (2). Telomerase synthesizes *de novo* telomeric repeats (TTAGGG) at chromosome ends using an integral RNA (termed TER or TR) and a telomerase reverse transcriptase (TERT) subunit that copies a short template sequence from the RNA component onto the telomere 3’end (3). Human TERT (hTERT) expression is suppressed in most human somatic tissues; consequently, telomeres gradually shorten with each cell division. When a few telomeres in a given cell become critically-short, they trigger a DNA damage signal and cellular senescence. Very short or otherwise unprotected telomeres can induce genomic instability, accumulation of mutations and a stress response, which are drivers of cancer and age-related diseases (4-6).

The large difference in telomere length between two closely related and inter-breedable species of mice, *Mus musculus* and *M. spretus*, was shown to be associated with a single locus encoding a DNA helicase, which was consequently termed ‘regulator of telomere elongation 1’ (RTEL1) (7). Indeed, RTEL1 was found as a dominant factor regulating telomere length and was suggested to facilitate telomere extension by telomerase in a yet unknown mechanism (7-10). RTEL1 is an essential DNA helicase from the XPD family helicases, which contain a conserved iron-sulfur cluster domain, translocate on ssDNA in a 5’-3’ direction, and play important roles in genome stability (11-13). In addition to its role in telomere length regulation, RTEL1 was suggested to play multiple roles in telomeric, as well as non-telomeric, genome stability: It associates with the replisome through binding to proliferating cell nuclear antigen (PCNA) to facilitate both genome-wide and telomeric replication (14); it is recruited to the telomeres by TRF1 to resolve G4 quadruplexes and prevent telomere fragility (15); it is recruited to telomeres by TRF2 in S phase to promote t-loop unwinding and prevent stabilization of reversed replication forks by telomerase (16-18); and it stabilizes long G-overhangs in cells overexpressing telomerase (19). In addition, RTEL1 is involved in non-coding RNA metabolism: it facilitates pre-U2 snRNA trafficking between the nucleus and cytoplasm (20) and the abundance and localization of the telomeric long non-coding RNA TERRA (21).

Heterozygous carriers of *RTEL1* mutations display decreased genomic stability and an elevated risk for cancer and pulmonary fibrosis (22-24). Biallelic *RTEL1* germline mutations cause a severe telomere biology disease (TBD) named Hoyeraal-Hreidarsson syndrome (HHS) (25-29) (reviewed in Hourvitz, 2023 #7843}). HHS is a severe form of dyskeratosis congenita, characterized by accelerated telomere shortening and diverse clinical symptoms including immunodeficiency, neurodevelopmental defects, and bone marrow failure (30). While accelerated telomere shortening is thought to be the main cause for the disease, the contribution of other manifestations of RTEL1 dysfunction, such as genome-wide DNA damage and chromosomal aberrations, to the diverse clinical symptoms remains to be explored (13).

We have previously found an HHS-causing mutation in a conserved methionine at position 492 of *RTEL1* (M492I) (27). The HHS patients were compound heterozygous for this mutation, *Rtel1*^M492I^, in combination with a nonsense mutation, *RTEL1*^R972X^, that leads to mRNA degradation by the nonsense-mediated decay pathway (27). Telomeres were severely shortened in leukocytes of the patients, but not in early passage primary fibroblasts (31). Yet, primary patient fibroblasts displayed diminished telomeric overhangs and increased DNA damage foci colocalizing with telomeres, suggesting that a defect in telomere structure, rather than length, compromised telomere protection (31). The ectopic expression of a 544 amino-acid long variant of TPP1 (TPP1-L), which does not facilitate telomere elongation, rescued the viability of patient fibroblasts expressing ectopic hTERT while their telomeres continued to shorten. Altogether, these observations suggested a defect in telomere structure, in addition to excessive telomere shortening in the blood, in these HHS patients (9,31).

A natural variation at the same position of RTEL1 in *M. spretus* (M492K) shortened the telomere length setpoint when introduced into *M. musculus*, generating a mouse strain with human-length telomeres termed ‘Telomouse’ (10). The structure and function of Telomouse telomeres appear normal, consistent with the natural presence of the K492 variation in *M. spretus* (10). Here, we derived an additional *Rtel1* mutant mouse strain, which is homozygous for the HHS-causing mutation (M492I), and termed it ‘HHS mouse’. We hypothesized that a mouse carrying the HHS-causing mutation in *Rtel1* would exhibit length-independent telomere dysfunction, genome-wide DNA damage and genomic instability, as found in HHS patients. Here, we report that the HHS mouse exhibits only moderate telomere shortening but more significant DNA damage at telomeres than observed in Telomice. These results indicate that the two *Rtel1* mutations affect different roles of RTEL1 in telomere length regulation and protection.

## MATERIALS AND METHODS

### Derivation of *Rtel1*^M492I^ mice

CRISPR guide RNAs were designed as described (32). For sgRNA preparation, the T7 promoter sequence was added to the sgRNA template by PCR amplification using pX335 (Addgene #42335) as a template with Phusion high-fidelity DNA Polymerase Kit (NEB), Common sgRNA-R: 5’-AGCACCGACTCGGTGCCACT-3’and Primer 1 with gRNA sequence underlined and template sequence in lower case: 5’-TTAATACGACTCACTATAGG<u>CATCTGCATCTCCAGAGCAA</u>gttttagagctagaaatagc-3’ or Primer 2 with gRNA sequence underlined and template sequence in lower case: 5’-TTAATACGACTCACTATAGG<u>CACCTGGAGGTCACAACACT</u>gttttagagctagaaatagc-3’ PCR products were purified using the QIAquick Gel Extraction kit (Qiagen), followed by *in vitro* transcription with the T7 High Yield RNA Synthesis kit (NEB). Newly synthesized sgRNAs were purified using the MEGAclear kit (Life Technologies), precipitated and diluted in injection buffer (10 mM Tris, mM EDTA, pH 7.5) at a concentration of 500 ng/μl. The quality of the sgRNAs was confirmed using the total RNA Nano Chip on an Agilent bioanalyzer. The final injection mix was prepared using ssDNA repair template (IDT; 100 ng/μl), sgRNA1 (50 ng/μl), sgRNA2 (50 ng/μl), and Cas9 Nickase mRNA (Trilink; 100 ng/μl) in injection buffer. The sequence of the ssDNA repair template is shown below. 5’-GTTCGTACCCTTATCCTCACCAGCGGTACCCTGGCTCCACTGTCTTCCTTTGCTCTGGAGATTCAGATGTATGTATGAGTCACCTGGAGGTCACAACACTAGGAACATGGTGGGTGGGGTTGG-3’ The final mix was spun twice at top speed for 30 minutes at 4°C to remove debris to avoid needle clogging. Cytoplasmic injection was done using C57BL6 zygotes. After injection, SCR7 (Xcessbio) was added to the overnight egg culture at the concentration of 50 μM. Out of the 15 mice born, one had the targeted allele.

### Preparation and culture of MEF

Embryonic day (E)13.5 mouse embryos were dissected with head and inner organs removed, rinsed in HBSS, then digested with 300 μl papain isolation enzyme (ThermoFisher) for 30 minutes at 37°C following the manufacturer’s protocol. Embryonic digests were transferred to conical tubes with 1 ml of HBSS, pipetted up and down to achieve single cell suspension and then spun. The pellets were resuspended in MEFs media (DMEM with 20% FCS, Pen/Strep/non-essential amino acids) and plated. MEFs were grown in DMEM media containing 2 mM L-glutamine, penicillin-streptomycin, non-essential amino acids and 20% fetal bovine serum until immortalization [around population doublings (PD) 60-70] and then with 10% fetal bovine serum in the same medium. BIOAMF-2 complete media was used for poorly growing homozygous *Rtel1*^M492I^ (‘I/I’ for short) cultures at early PD, then 20% fetal bovine serum was used. All media and media supplements were purchased from Biological Industries Inc., Beit Haemek, Israel. Four mouse embryonic fibroblast cultures were grown over 250 PD and sampled at different PD for the different experiments.

### Culture of S2 human fibroblasts

Fibroblasts derived from HHS-patients were grown in DMEM media supplemented with 10% fetal bovine serum, penicillin-streptomycin, 2mM L-glutamine and non-essential amino acids. To induce the ectopic expression of human RTEL1 fron a tetracycline-inducible (TET-on) promoter (9), the medium was supplemented with 30 ng/ml doxycycline (Dox). To silence its expression, Dox was omitted from the medium and a certified tetracycline-free fetal bovine serum (04-005-1A, Biological industries) was used. Media and media supplements were purchased from Biological Industries Inc. Beit Haemek, Israel.

### Genomic DNA extraction

Leukocytes were obtained from mouse blood by lysing red blood cells in RBC lysis buffer (155 mM NH_4_Cl, 10 mM KHCO_3_ and 0.1 mM EDTA pH 8) followed by centrifugation. Mouse leukocytes, MEFs, and S2 human fibroblasts were lysed in 10 mM Tris pH 7.5, 10 mM EDTA, 0.15 M NaCl, 0.5% SDS and 100 μg/ml proteinase K overnight at 37°C. Mouse tail samples were lysed in 100 mM Tris HCl pH 8.5, 5 mM EDTA, 0.1% SDS, 200 mM NaCl and 100 μg/ml proteinase K overnight at 50°C. Following cell lysis, high molecular weight genomic DNA was phenol-extracted, ethanol-precipitated, and dissolved in TE. Genomic DNA samples were examined by agarose gel electrophoresis of undigested DNA, and degraded samples were excluded from further analysis. DNA sequenced by *NanoTelSeq* was extracted either by the standard proteinase K/phenol method or by Monarch genomic DNA purification kit (NEB #T3010).

### In-gel analysis of telomeric restriction fragments

Genomic DNA was treated with RNase A, digested with the *HinfI* restriction endonuclease, and quantified by Qubit fluorometer. 1-5 μg (equal amounts in each gel) of the digested DNA samples were separated by pulsed-field gel electrophoresis (PFGE) using a Bio-Rad CHEF DR-II apparatus in 1% agarose and 0.5 x TBE, at 14°C, 200V and pulse frequency gradient of 1 sec (initial) to 6 sec (final) for 18-22 hr, or in regular gel apparatus, 0.7% agarose and 1 x TBE for 1,300 V x hr. Gels were ethidium-stained, imaged, dried under vacuum for 1 hr at room temp and then 1 hr at 50°C. DNA was denatured by incubating the dried gel in 0.5 M NaOH and 1.5 M NaCl for 30 min, neutralized in 1.5 M NaCl and 0.5 M Tris–HCl pH 7.0 for 30 min, rinsed with H_2_O, and hybridized as previously described (10). Probe mixtures contained a telomeric oligonucleotide (AACCCT)3, and mid-range PFG ladder (NEB Inc.; for mouse samples) and 1 kb ladder (GeneDirex Inc.; for human DNA samples), which were digested with *HinfI*, dephosphorylated by quick calf intestinal phosphatase (NEB Inc.) and heat inactivated. All probes were 5’ end-labeled with [alpha-32P]-ATP and T4 polynucleotide kinase (NEB Inc.). The gels were visualized by PhosphorImager. TRF length was quantified using *TeloTool* (33) (corrected mode). The results for the MEF cultures and mouse samoples are detailed in Supplementary Table S1 and S3, respectively.

### 2D gel hybridization

Genomic DNA was treated with RNase A, digested with *HinfI* restriction endonuclease, and quantified by Qubit fluorometer. In the first dimension, equal amounts (∼ 20-25 μg) of the digested DNA samples were separated by PFGE using a Bio-Rad CHEF DR-II apparatus, in 0.4% agarose and 0.5 x TBE, at 20°C, 35V and pulse frequency gradient of 1 sec (initial) to 6 sec (final) for 50 hr. The gel was stained with ethidium bromide and visualized by UV light. The appropriate lanes were then excised for the second-dimension PFGE in a 1.1% agarose gel, 0.5 x TBE, and 0.3 μg/ml ethidium bromide, which was poured around the excised slab. The second-dimension electrophoresis was carried out in the same apparatus at 14°C, 200V and pulse frequency gradient of 1 sec (initial) to 6 sec (final) for 20-22 hr. The gel was then visualized using UV light, dried, hybridized as described above, and visualized by PhosphorImager.

### Restriction fragment length PCR (RFLP) to verify MEFs genotypes

PCR reaction was performed on gDNA (50-100 ng) to amplify a 269 bp product, using the following primers in 25μl reaction (annealing Tm 58; extension 72’ 15’’; 35 cycles):

> 1 – mRtel1gM492UP 5’-CAGTCCCAGCCAGAGTATGC-3’
>
> 2- mRtel1gM492LP 5’-AATGGACTGGGAATGGCCAG-3’

The products were cut by *HinfI* restriction enzyme, giving the following results according to the genotypes: M/M: 117+152 bp, M/I:100+117+152 bp, I/I: 100+17+152 bp

### Western Blot

Western analysis was performed as previously described (9), but using alpha tubulin antibodies as a loading control.

### Immunofluorescence of cultured cells

Cells were seeded onto glass coverslips and grown for 1-4 days. Immunofluorescence was done as described (9). Imaging was performed using a FV-1200 confocal microscope (Olympus, Japan) with a 60X/1.42 oil immersion objective. NIH ImageJ28 was used for image analysis and foci quantification.

### Meta-TIF assay

Meta-TIF was performed as previously described (10), and images were acquired using the Olympus IX83 fluorescence microscope system (Olympus).

### Metaphase FISH

Metaphase spreads and FISH were performed as previously described (10), using TelC-Alexa488 telomeric probe (F1004) or TelC-Cy3 probe (F1002-5; Panagene Inc). Imaging acquisition was done with an Olympus IX83 fluorescence microscope system.

### Chromosome-Orientation Fluorescent In-Situ Hybridization (CO-FISH)

The protocol was performed as previously described (10), and images were acquired using the Olympus IX83 fluorescence microscope system (Olympus).

### Nanopore sequencing of telomeres

Three or Five samples were pooled in a sequencing library using barcoded telorette 3 oligonucleotides (10). The barcoded telorette oligonucleotides were first annealed to a complementary teltail-tether oligonucleotide and then each double-stranded barcoded telorette was ligated to an individual DNA sample (1 - 4 μg each). The ligation reactions were stopped by adding 20 μM EDTA and the barcoded samples were combined, purified, ligated to sequencing adapter (AMII) and purified again. 1-5 μg of the sequencing library was loaded onto a Nanopore R9.4.1 MinION flow cell following the standard protocol for native barcoding ligation (SQK-LSK109 with EXP-NBD104; Oxford Nanopore Technologies). Oligonucleotides used for sequencing:

### Telorette3-NB01n

P-5’-TGCTCCGTGCATCTCC-AAGGTTAA-CACAAAGACACCGACAACTTTCTT-CAGCACCTCTAACC-3’

### Telorette3-NB02n

P-5’-TGCTCCGTGCATCTCC-AAGGTTAA-ACAGACGACTACAAACGGAATCGA-CAGCACCTCTAACC-3’

### Telorette3-NB03n

P-5’-TGCTCCGTGCATCTCC-AAGGTTAA-CCTGGTAACTGGGACACAAGACTC-CAGCACCTCTAACC-3’

### Telorette3-NB04n

P-5’-TGCTCCGTGCATCTCC-AAGGTTAA-TAGGGAAACACGATAGAATCCGAA-CAGCACCTCTAACC-3’

### Telorette3-NB05n

P-5’-TG CTCCGTGCATCTCCAAGGTTAAAAGGTTACACAAACCCTGGACAAGCAGCACCTCTAACC-3’

### Teltail-tether

5’-AACCTTGGAGATGCACGGAGCAAGCAAT-3’ or 5’-TTAACCTTGGAGATGCACGGAGCAAGCAAT-3’

### Computational processing of Nanopore sequencing reads

Deducing the nucleotide sequence (basecalling) from the raw data (Pod5 files) was performed with the Nanopore Dorado basecaller (versions 0.3.0 - 0.6.2) in the super-accurate mode. The sequences (FastQ files) were filtered for telomere reads by *TelomereAnalyzer* <u>version 1.03 or</u> 1.0.8-beta, searching for the telomeric repeat PyPyAGGG (Py=C or T) in the reverse-complement read sequences while allowing one mismatch in each 6 nt repeat (34). The telomere reads were presented as plots of telomere density (0 to 1; with and without 1 mismatch) and the telomere length was calculated as previously described from the telomere density allowing mismatched repeats (10).

### Generating a three-dimensional structure model of the murine and human RTEL1 helicase domain

There are no available structures of the full length RTEL1 from any organism. The only structures of RTEL1 currently available publicly are those of the Harmonin-homology domain 2 (HHD2) of human RTEL1 (RCSB PDB ID: 7WU8) and the full-length model generated by the artificial intelligence software *AlphaFold* (35) (human AF-Q9NZ71-F1 and Mouse AF-Q0VGM9-F1). Structural comparison of the Alphafold generated RTEL1 helicase domain with that generated by the *Protein Homology Recognition Engine Phyre 2* (10,36) shows striking similarity between the two structures; therefore, we used the *AlphaFold* structure of the helicase domain of human and mouse RTEL1 for our analysis. Similarly to our previous RTEL1^M492K^ study, the positioning of the single stranded DNA at the RTEL1 helicase domain was predicted using an overlay of the *AlphaFold* RTEL1 model with its closest structural homolog, the crystal structure of the DinG XPD family helicase in complex with single stranded DNA (ssDNA; RCSB PDB ID: 6FWS (11)). The *AlphaFold* Helicase domain of human and mouse RTEL1 bound to ssDNA was used for any subsequent analysis.

### Statistical analysis

Statistical analysis was performed by Microsoft Excel and GraphPad Prism 8.0, using simple linear, polynomial, or segmental linear regression with setting the break points manually to give the best fit by coefficient of determination (R2). For statistical tests two-tailed paired or unpaired t-test were used, as indicated for each figure. P < 0.05 was considered statistically significant.

## RESULTS

### Derivation of the HHS mouse model

The sequence of the RTEL1 helicase domain is highly conserved between human and mouse (Figures 1A, B and Supplementary Figure S1). Specifically, methionine 492 is conserved in vertebrates with only a few naturally occurring exceptions: a leucine found in this position in several cat species, and a lysine in *M. spretus* (10,27). To test the effect of the M492I HHS-causing mutation in the mouse, we derived a *M. musculus* strain with this mutation using CRISPR-Cas9 assisted genome editing in zygotes (Figure 1C). We employed the Cas9 nickase (D10A Cas9 mutant) enzyme and two guide RNAs to generate nicks on both strands of the *Rtel1* gene to facilitate gene conversion using a repair template (Figure 1C). Successful gene targeting in the resulting offspring was confirmed by DNA sequencing (Figure 1D). This founder mouse was backcrossed to the isogenic parental wild-type (WT) C57BL/6 mice to generate F0 mice heterozygous for the *Rtel1*^M492I^ allele, termed *Rtel1*^M/I^. F0 *Rtel1*^M/I^ mice were crossed to C57BL/6 mice two additional times to generate F1 and F2 *Rtel1*^M/I^ mice. Homozygosity for the *Rtel1* mutation (*Rtel1*^I/I^) was established in the third generation (F3). The homozygous *Rtel1*^I/I^ (I/I for short) mice were then bred *inter se* for eleven additional generations up to F14 to date. We termed the *Rtel1*^I/I^ mouse model ‘HHS mouse’.

**Figure 1.**
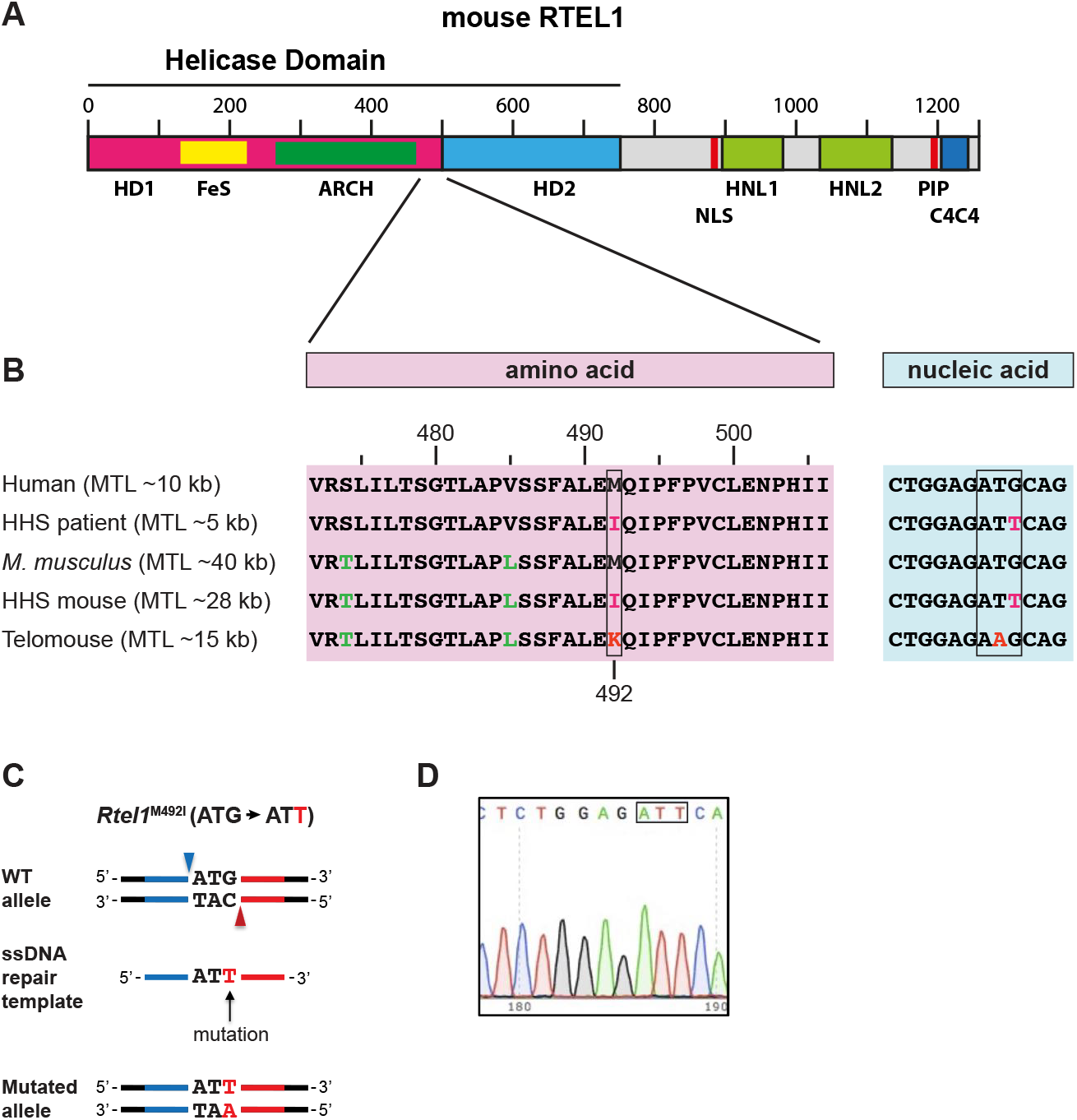
Derivation of the *Rtel1*^M492I^ allele by CRISPR-Cas9 nickase editing. (A) A scheme showing the helicase domains (HD1 and HD2), the iron-sulfur cluster (FeS), the nuclear localization signal (NLS), harmonin N-like domains (HNL1 and HNL2), PCNA-interacting protein (PIP) motif, and the C4C4 ring-finger domain (13) of RTEL1. (B) RTEL1 protein and DNA sequence alignment showing codon 492 (boxed) and flanking sequences in human and mouse. Variations in mouse from the human sequence are shown in green and mutations in HHS patients and mouse models are shown in red. (C) Illustration showing the WT allele, the two positions targeted by gRNAs (blue and red arrowheads), the ssDNA repair template with the mutation, left homology arm (blue), right homology arm (red), and the resulting mutant allele. (D) Sanger sequencing of the HHS mouse founder line confirming the replacement of the ‘G’ with a ‘T’ in codon 492, changing the methionine to an isoleucine.

### The HHS mutation causes telomere shortening in mouse embryonic fibroblasts and tissues

Telomerase positive HHS patient cells such as EBV transformed lymphoblastoid cell lines (LCLs) and primary fibroblasts expressing ectopic hTERT display progressive telomere shortening despite the presence of catalytically-active telomerase (9,27,31). To examine if the *Rtel1*^M492I^ mutation has a similar effect on telomere length in the mouse, we established mouse embryonic fibroblasts (MEFs) from heterozygous (M/I), homozygous (I/I) and (M/M) F3 littermate embryos generated by intercrossing F2 *Rtel1*^M/I^ mice. The WT MEFs might have inherited short telomeres from their heterozygous parents and therefore are indicated as M/M* to distinguish them from WT (M/M) MEFs generated by breeding WT mice that did not harbor the *Rtel1*^M492I^ mutation in their pedigree ancestors, avoiding trans-generational inheritance of short telomeres. MEF cultures were established by serial passaging and selection for spontaneous immortalization, and followed in culture over 250 population doublings (PD). The genotypes of the MEFs were reconfirmed at PD 250 by PCR and differential digestion of the *Rtel1*^M492I^ allele by *HinfI* (Figure S2). During growth, the cultures passed several crisis periods and one to three transition points in which the MEFs increased their growth rates and changed their cellular morphology (Figure S3). All MEF lines were immortalized by PD 70 and their shapes became rounder. During the final growth phase, the M/I, I/I, M/M* and M/M MEF cultures grew at the same average rate, about one PD/day (Figure S3A, B). We examined the TRF length for these four cultures at multiple PD levels by in-gel hybridization (Figure 2A). Each sample was assayed several times in independent gels and the mean TRF length (MTL) was calculated (Figure 2B and Supplementary Table S1). The average MTL of all MEFs cultures shortened during the initial growth phase until immortalization occurred at about PD 70 (arrows in Figure 2A, B), presumably due to the lower telomerase activity present in MEFs prior to immortalization (10,37). After immortalization, MTL of the WT and M/I cultures stabilized, while the I/I MTL continued to shorten at an average rate of ∼35 bp per PD to 23.7 kb at PD 250, as compared with the WT (M/M) MTL of 37.8 kb (Figure 2A, B). The M/M* MEFs had shorter MEFs MTL of 29.7 kb at PD 10, presumably due to the inheritance of short telomeres from the heterozygous parents. Altogether, the *Rtel1*^M492I^ mutation caused gradual telomere shortening in mouse fibroblasts, phenocopying the telomere shortening observed in human HHS patients carrying the same mutation (Figure S4 and (9)). However, the telomeres in the I/I MEFs did not shorten as much as in the K/K MEFs, reaching a MTL of 23.7 kb at PD 250, as compared to 14.7 kb, respectively (Figure 2B and (10)). Consistently, also the telomeres of the heterozygous M/I MEFs shortened significantly less than those of the M/K MEFs (Figure 2B and (10)). Next, we asked whether the *Rtel1*^M492I^ mutation caused a significant accumulation of critically short telomeres, which are the most relevant to physiology and cell fate. To this end, we sequenced individual telomeres by *NanoTelSeq* (10). Consistent with the TRF analysis, at PD 250, the median telomere length measured by *NanoTelSeq* in the Rtel1 I/I MEFs was 23.7 kb, as compared to 5.4 and 29.2 kb for Rtel1 K/K and Rtel1 M/M MEFs, respectively (Figure 2C and Supplementary Table S2). We did not identify any telomere shorter than 5 kb in the I/I MEFs, rulling out increased accumulation of critically-short telomeres.

**Figure 2.**
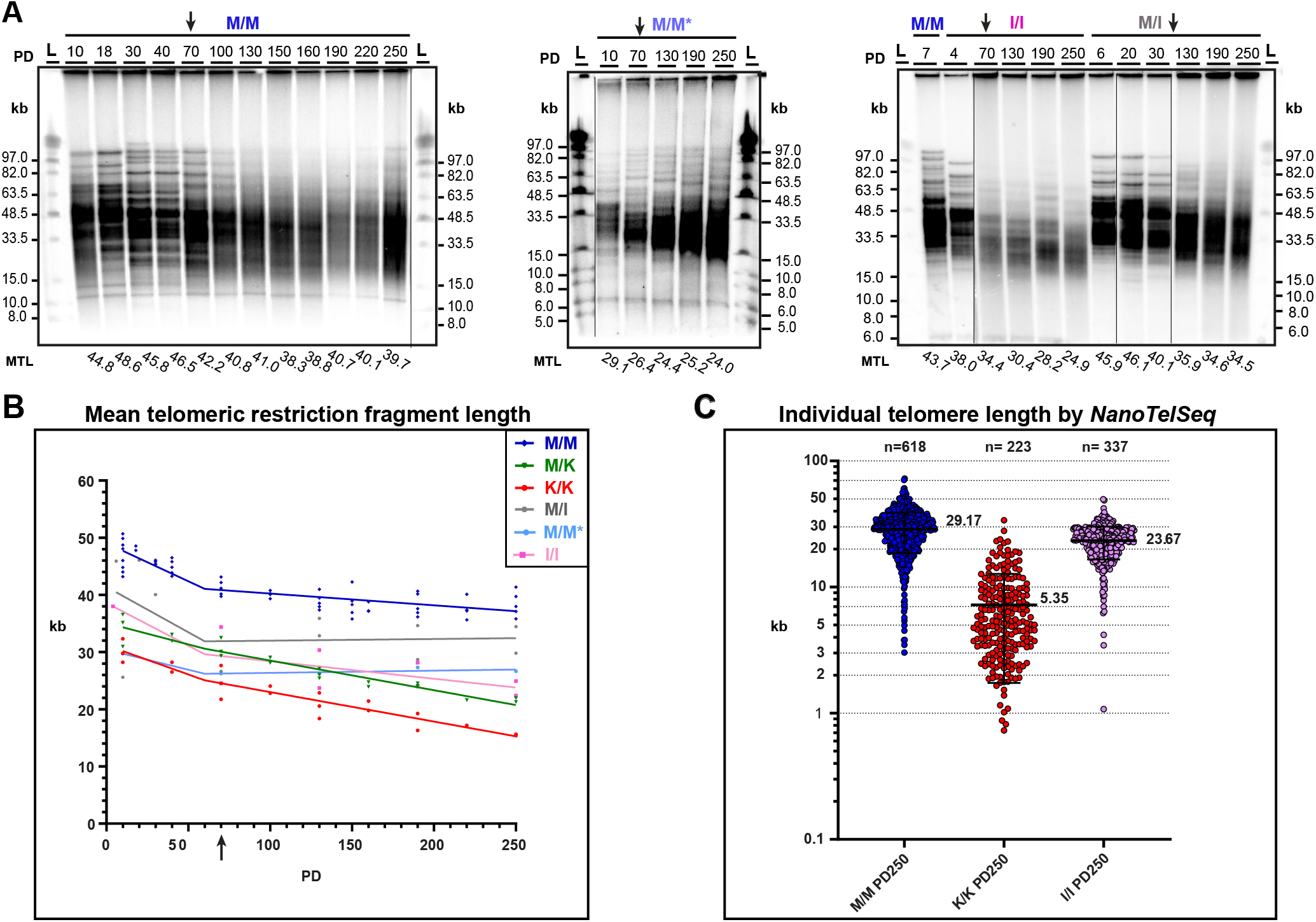
Telomeres of MEFs carrying the homozygous *Rtel1*^M492I^ mutation progressively shorten with cell division in culture. MEFs cultures were prepared from F3 littermate embryos, I/I, M/I and M/M*, generated by mating F2 heterozygous mice carrying the Rtel1 M492I mutation. Another control M/M MEFs culture (M/M) was generated by intercrossing WT mice which did not have any Rtel1 mutation in their pedigree, to avoid trans-generational inheritance of short telomeres from heterozygous *Rtel1* mutant mice to the WT offspring. All four cell lines were immortalized by serial passaging and grown to PD 250. (A) Genomic DNA samples extracted from the indicated MEFs cultures and PDs were digested by *HinfI* restriction endonuclease and analyzed by PFGE, denaturation and in-gel hybridization with a C-rich telomeric probe.. Mean TRF length (MTL) was quantified for each sample by *TeloTool* (33) and indicated below the lanes. (B) MTL values measured in multiple gels for the same M/M, I/I, and M/I MEFs samples (Supplementary Figure S1). For comparison, MTL values for K/K and M/K MEFs were imported from (10). (C) Telomeres from the indicated cultures at PD250 were sequenced by *NanoTelSeq* (I/I) or re-basecalled and processed from the raw data of previous sequencing reactions (M/M and K/K) reported in (10). Scatter plots show the individual telomere length of the indicated MEF cultures. Mean and standard deviation (SD) are indicated by horizontal lines. Median values in kb are indicated to the right of each scatter plot, and “n” indicates the number of telomeric reads. The details of all individual telomere reads are shown in Supplementary Table S2.

The MTL of the I/I MEFs shortened at an average rate of ∼35 bp per PD. To examine the rate of telomere shortening caused by the the *Rtel1*^M492I^ mutation in human cells, we employed telomerase-positive HHS patient fibroblasts that elongated their telomeres and maintained growth upon inducible ectopic expression of WT RTEL1v2 (S2 in (9); Figure S4). Omitting doxycycline from the medium silenced the ectopic RTEL1v2, caused gradual telomere shortening over 100 PDs, slowed down cell growth after 60 PD, and changed cell morphology (Figure S4). While the MTL was stable in the cells expressing RTEL1v2 (DOX+), it shortened gradually upon silencing RTEL1v2 from 12.3 kb at PD 10 to 6.4 kb at PD100 (Figure S4B), indicating an average shortening rate of 65 nt per PD, about twice the rate of shortening observed in I/I MEFs. Sequencing of individual telomeres by *NanoTelSeq* revealed that the median telomere length shortened from 10.2 kb at PD 10 to 3.7 kb at PD 100, indicating an average shortening rate of 72 bp. At PD 100 without RTEL1v2, 19% and 4% of the telomeres were shorter than 3 kb and 2 kb, respectively, while no such short telomeres were found at PD 10 or PD 50. These results suggest that telomeres shorter than 3 kb or 2 kb are responsible for slowing cell growth beyond PD 60 (Figure S4A).

To examine the effect of the *Rtel1*^M492I^ mutation on somatic tissues of HHS mice, we measured the telomere length in blood and tail DNA samples over several generations (F3 to F10 in blood and F3 to F11 in tail) and compared the results to Telomice (K/K) and WT (M/M) control mice (Figure 3). In blood, the average MTL shortened to an average length of 27.9 kb in F8 HHS mice and no further shortening was apparent through F10, as compared to 40.1 kb in WT samples (Supplementary Table S3 and Figures 3A, C and S5). In tail samples, MTL shortened to an average of 29.0 kb in F6 HHS mice and no further shortening was observed through F11, as compared to 42.3 kb in the WT samples (Supplementary Table S3 and Figures 7B, D and S5). These results indicate that the telomeres in the HHS mouse germline had stabilized at a new telomere length set point that is significantly longer than the new setpoint reached in Telomouse F16 (16.7 kb and 19.2 kb in blood and tail, respectively; Figure 3C, D) and (10). Moreover, measuring the length of individual telomeres in the blood and tail of an F14 HHS mouse, F14 Telomouse, and WT mouse by *NanoTelSeq* revealed median telomere lengths of 19.5 kb and 17.8 kb in the HHS mouse blood and tail respectively, compared to 7.0 and 6.4 kb in Telomouse and 35.2 kb in the WT mouse blood (Figure 3E). We did not observe a significant elevation in telomeres shorter than 3 or 2 kb in the HHS mouse, compared to Telomouse.

**Figure 3.**
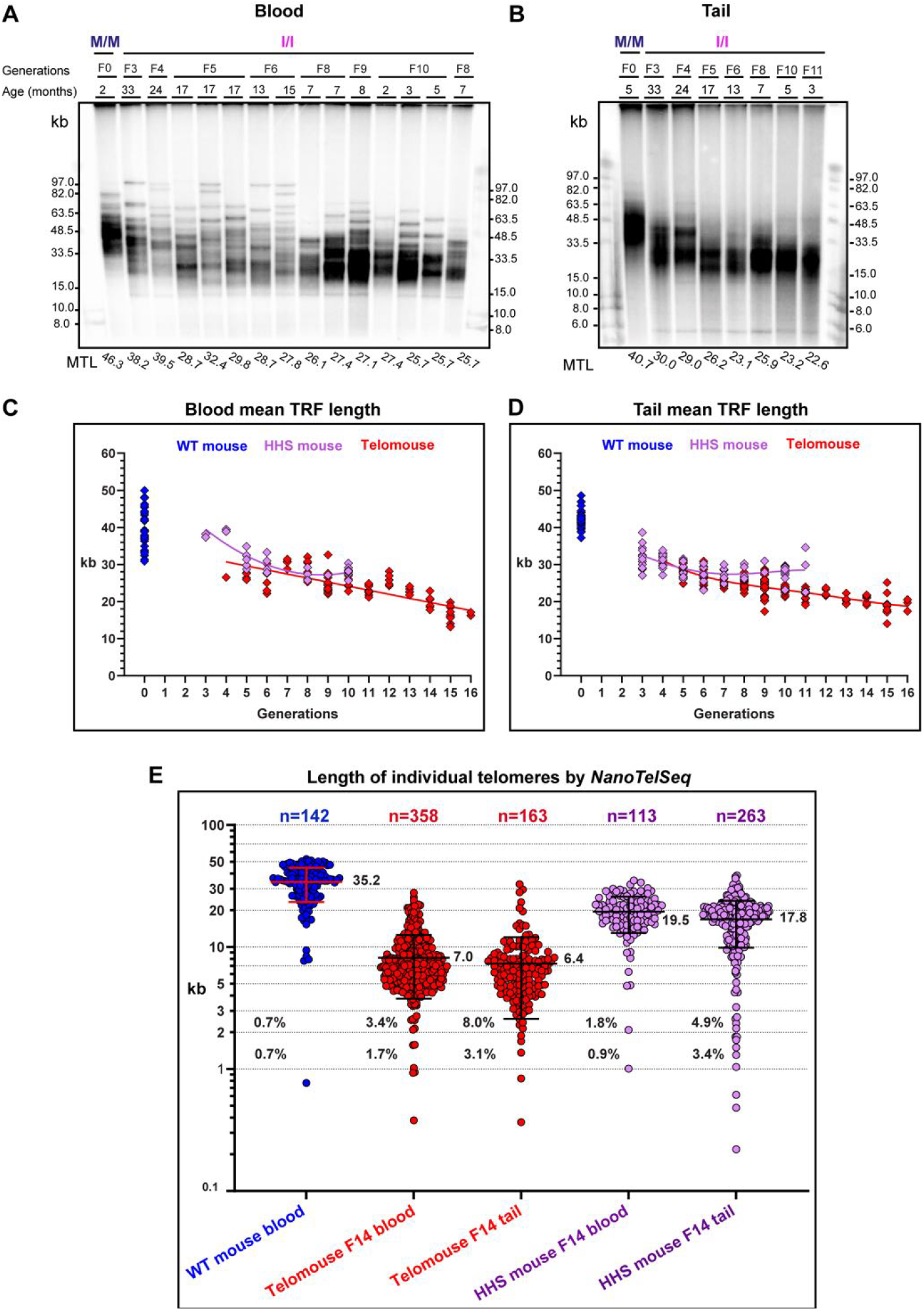
HHS mouse telomeres progressively shortened over generations but less than Telomice telomeres. Genomic DNA samples extracted from blood leukocytes (A) or tail (B) of HHS or WT mice at the indicated generations and ages were analyzed by PFGE and in-gel hybridization to the denatured DNA. Representative gels are shown here and additional gels in Figure S5. All MTL values measured were plotted for the blood (C) and tail (D) samples. Data for WT mice and Telomice were adapted from (10). The best-fit regression lines are shown (2^nd^ order polynomial for blood and 4^th^ order polynomial for tail). The average ages of the mice sampled were 329 days for HHS mice, 331 days for Telomice, and 355 days for WT mice. All HHS mouse data are summarized in Supplementary Table S3. (E) Scatter plots show the length of individual telomeres in the indicated mouse samples measured by *NanoTelSeq*. WT mouse (blue), Telomouse (red), and HHS mouse (purple) were all 13 months old. Mean and SD are indicated by horizontal lines. Median values in kb are indicated to the right of each plot and the percentage of telomeres below 3 and 2 kb to the left of the plots. ‘n’ indicates the number of telomeric reads. The details of all individual telomere reads are summarized in Supplementary Table S2.

Altogether, the *Rtel1*^M492I^ mutation caused gradual telomere shortening over 100 and 250 PD in both human and mouse fibroblasts, and over several generations of the HHS mouse. In the mouse, however, the *Rtel1*^M492I^ mutation caused milder telomere shortening than the *Rtel1*^M492K^ mutation, consistent with the more conservative nature of the substitution of methionine to isoleucine (both comprised of a hydrophobic side chain) than to the positively charged lysine.

### The HHS mutation impairs telomere protection and genome stability

The essential role of telomeres is to suppress the DNA damage response (DDR) at chromosome ends and prevent the consequent cell cycle arrest and deleterious fusions of chromosomes (1). Telomere dysfunction can occur either as a result of telomere shortening beyond a critical length or via damage to the telomere structure. DDR foci at telomeres, termed telomere dysfunction-induced foci (TIF), are considered a hallmark of telomere failure. In fibroblasts derived from HHS patients, the *Rtel1*^M492I^ mutation was associated with activation of DDR at telomeres even without telomere shortening, suggesting the presence of a length-independent defect in telomere structure and function (31). As shown above (Figure 2), in the MEFs, the HHS mutation *Rtel1*^M492I^ resulted in milder telomere shortening than the *Rtel1*^M492K^ mutation, raising the question whether mouse cells harboring the HHS mutation exhibit DNA damage independent of critically-short telomeres. We studied the presence of telomeric and genome-wide DDR foci in the I/I MEFs using antibodies against the DDR marker γH2AX and the telomere protein TRF2, and compared them to K/K and M/M (WT) MEFs (Figure 4). The overall (genome-wide) levels of DDR were high in all MEF cultures regardless of genotype (Figure 4A, B), as shown previously for MEFs growing under atmospheric oxygen levels (10,38). However, there was no significant difference in the abundance of DDR foci between the I/I MEFs and the K/K or M/M MEFs (Figure 4B). Some of the cells displayed overt damage throughout the nucleus without visible distinct foci, but these cells were present in all cultures (Figure 4D, E). DNA damage at telomeres, however, was significantly elevated in the I/I but not K/K MEFs, as revealed by the increase in the number of TIF (Figure 4C). Also elevated were the levels of micronuclei and anaphase bridges, which are known to associate with telomere dysfunction and genome instability (Figure 4D, E) (5). Altogether, these results suggest that the I/I MEFs telomeres, although longer than those of K/K MEFs, were nevertheless deprotected and compromised in their ability to suppress the DDR and prevent genomic instability.

**Figure 4.**
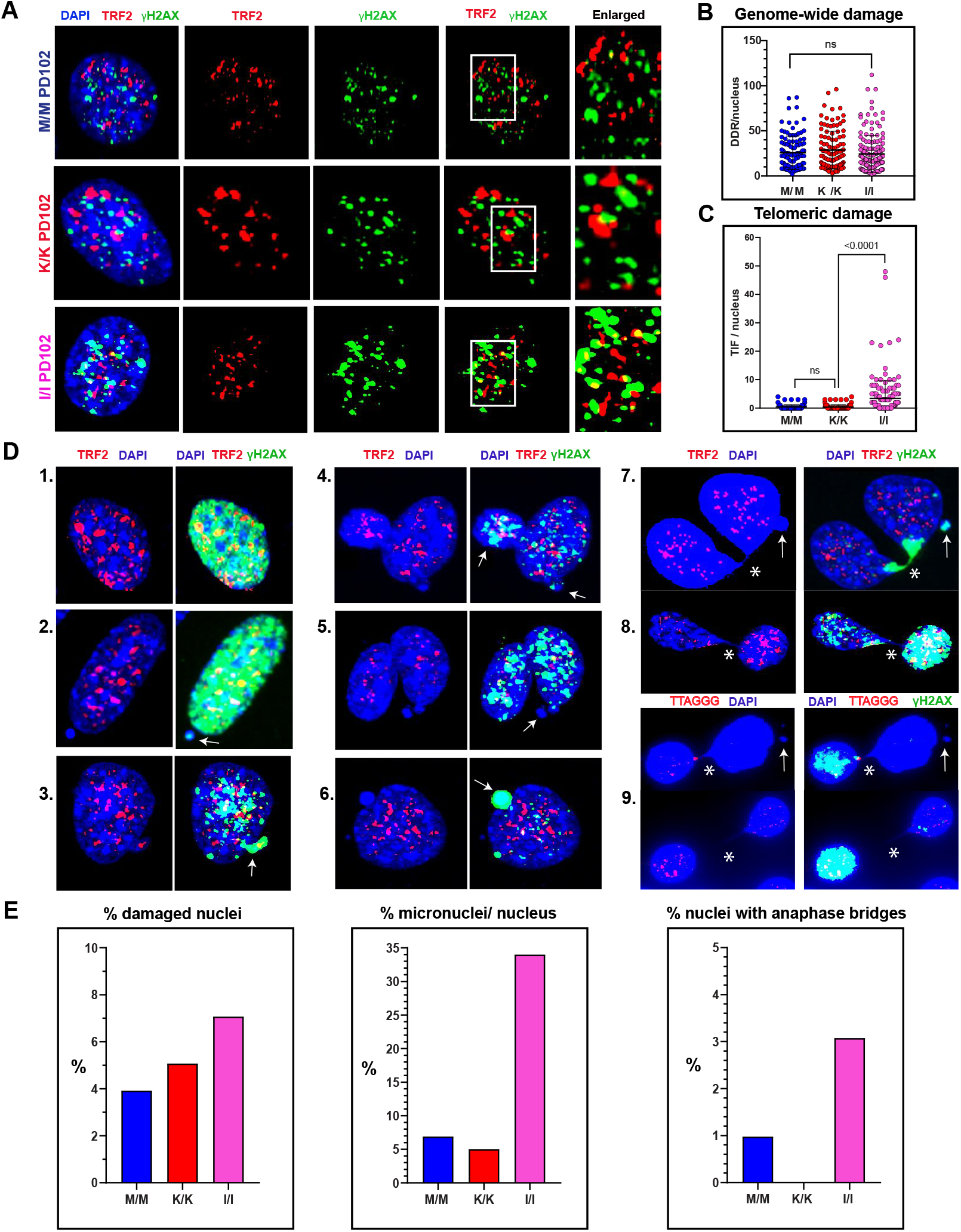
The HHS mutation impairs telomere protection and genome stability. (A) The formation of DNA damage response foci and their localization to telomeres (defined as TIF) in interphase nuclei were examined by immunofluorescence staining with antibodies for the DDR marker γH2AX (green) and the telomere protein TRF2 (red). Scatter plots show the number of DDR foci (B) and TIF (C) per nucleus. A total of 110 M/M, 110 K/K, and 153 (B) or 182 (C) of I/I nuclei were countified for MEFs PD 102. All three MEFs cultures were analyzed side by side; the data for the K/K and control (M/M) MEFs is adapted from (10). The mean and SD values are indicated by black lines, and P values were calculated by unpaired t-test. (D) Representative images showing nuclei with overt DDR (D1,2), micronuclei (D2-7 arrows), and anaphase bridges (D7-9 stars). (E) Bar graphs showing the percentage of nuclei with overt DDR, micronuclei, and anaphase bridges in a total number of 195-250 nuclei quantified for each genotype.

Since in interphase cells we can only observe telomeres with sufficient length and detectable telomeric signal, we studied TIF formation also in metaphase-arrested cells where individual chromosomes and their ends can be visualized by DAPI staining. Telomeric repeats were detected by FISH with a telomeric PNA probe, and DDR foci by immunofluorescence with γH2AX antibodies (Figure 5). Analyzing TIF in metaphase chromosomes of the I/I MEFs revealed that most of them were found in chromosome ends with detectable FISH signal and not at signal-free chromosome ends (Figure 5A). In addition, among the I/I MEFs we observed an increased level of anaphase bridges, in which at least one of the two bridged nuclei displayed overt DDR (Figure 5B). Since telomeres even shorter than the FISH detection limit did not apparently activate DDR in the K/K MEFs (10), the presence of TIF at longer telomeres with detectable FISH signal in the I/I MEFs suggested that the activation of DDR at these telomeres was driven by a defect in telomere structure or composition rather than insufficient length. RTEL1 dysfunction in patient cells and mouse *Rtel1* null cells was reported to induce telomere aberrations, such as telomere loss, telomere fragility, interstitial telomere insertion (ITS) and telomere fusion (reviewed in (13)). To examine the frequency of telomere aberrations caused by the HHS *Rtel1*^M492I^ mutation, we performed metaphase FISH on MEFs cultures PD 250 (Figure 6). The most significantly elevated phenotype in the I/I MEFs was telomere fragility (Figure 6B, C), consistent with the increased frequency of TIF in these MEFs (Figures 4C, 5A) and pointing to increased replication stress at telomeres. The I/I MEFs displayed additional phenotypes associated with genomic instability such as anaphase bridges, connected nuclei, micronuclei, chromosome fragments, and mitotic catastrophe, which could be the consequences of telomeric DNA damage (Figures 3D, E and 6D-F).

**Figure 5.**
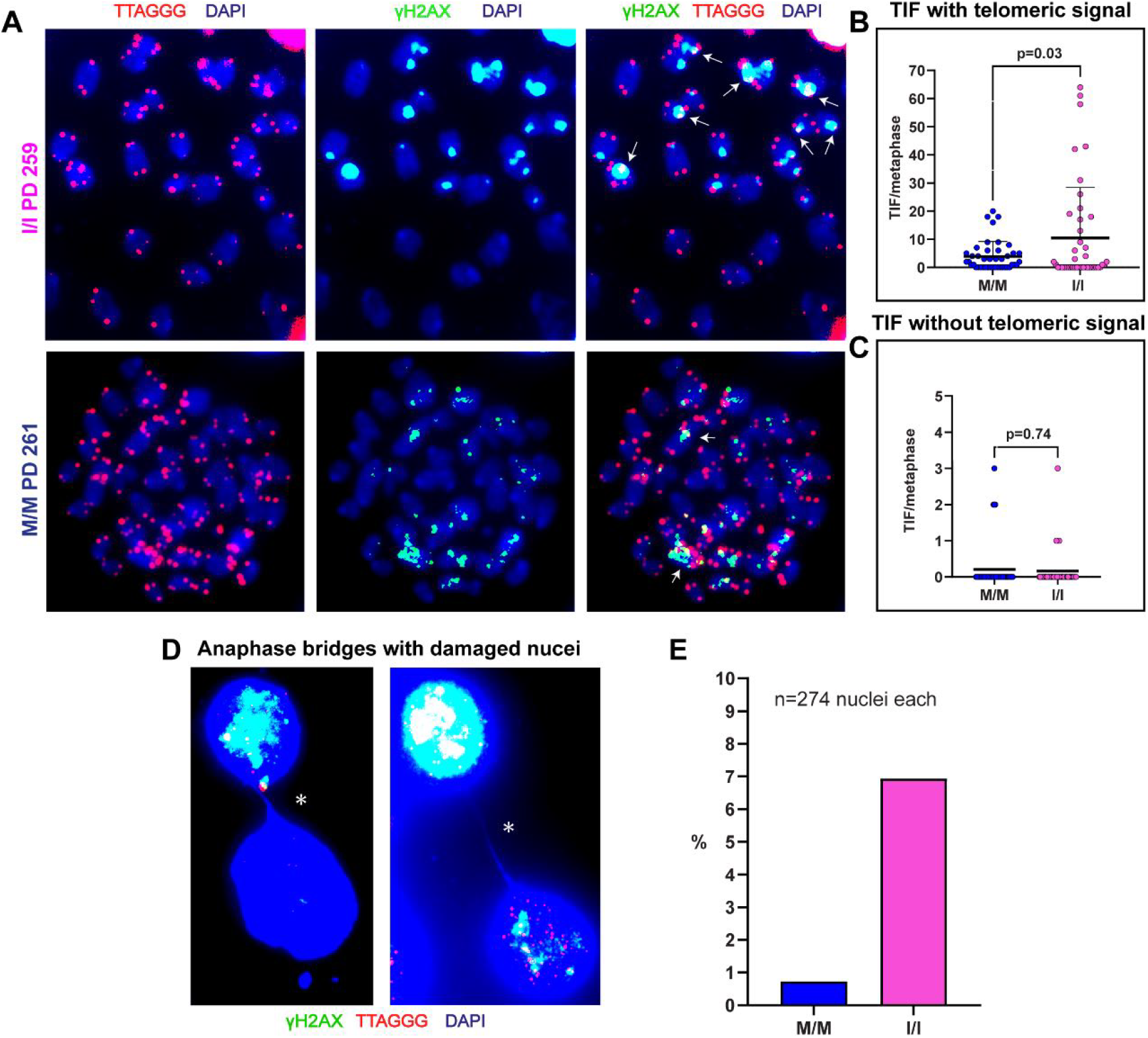
Telomeric DDR occurs at chromosome ends with detectable telomeric signal. (A) Metaphase TIF (indicated by white arrows) were analyzed by FISH-IF using a γH2AX (green) antibody and a telomeric C-probe (red). A total of 43 metaphases were countified for each M/M (PD 261) and I/I (PD 259) MEFs culture. Shown are the number of TIF per metaphase at chromosome ends with (B) or without telomeric signal (C). The mean and SD for TIF are indicated by black lines. P values were calculated by two-tailed paired t-test. (D) Examples of abundant pairs of I/I MEFs nuclei that are connected by anaphase bridges and at least one nucleus of the pair displays overt DDR. The DAPI signal is enhanced to visualize the bridge. A total of 274 nuclei were counted for each M/M (PD 261) and I/I (PD 259) MEFs culture. (E) Bar graphs showing the percentage of nuclei with anaphase bridges and overt DDR (in one or two of the bridged nuclei).

**Figure 6.**
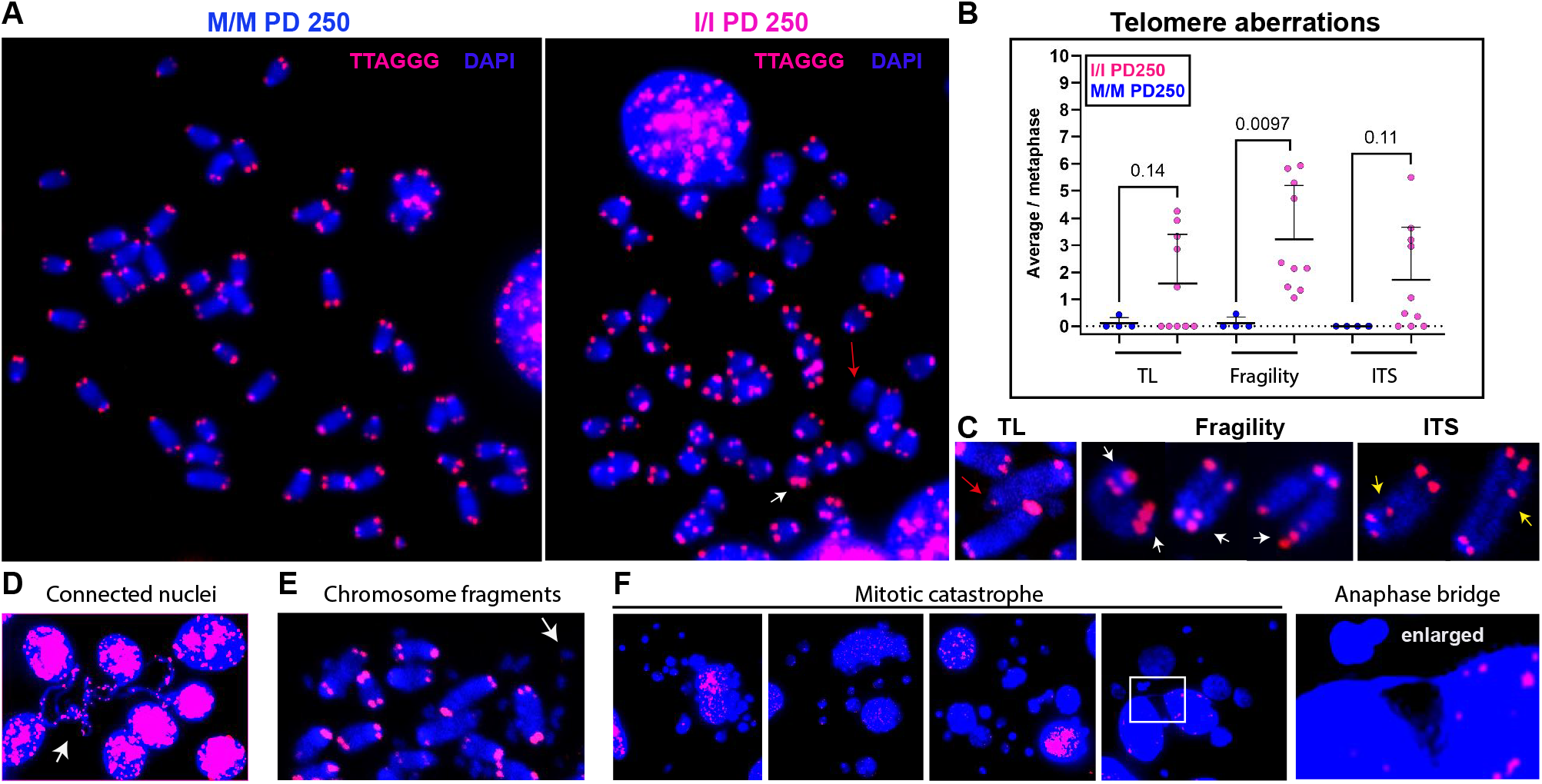
Increased telomeric and genomic aberrations in HHS MEFs. (A) Representative images for metaphase chromosomes of M/M and I/I MEFs at PD 250, hybridized with a telomeric probe (red). Colored arrows indicate chromosomal aberrations as in (C). (B) Scatter plots showing the average number of each aberration type per metaphase (normalized to the number of chromosomes counted). Horizontal lines show mean and SD. 619 chromosomes (15 metaphases) M/M PD250 and 692 chromosomes (17 metaphases) I/I PD 250 were quantified. P-values were calculated by unpaired two-tailed t-test. (C) Representative images of aberration types in I/I MEFs are shown: telomere loss (TL), telomere fragility, and interstitial telomere insertions (ITS). (D-F) Genomic abnormalities associated with the M492I mutation, including connected nuclei, chromosome fragments, mitotic catastrophe, micronuclei and anaphase bridges.

Telomere sister chromatid exchange (T-SCE), indicating elevated homologous recombination events between sister telomeres, was reported in RTEL1-deficient HHS patient cells (28). We examined the frequency of T-SCE events in the I/I MEFs by chromatid-orientation FISH (CO-FISH), differentiating between nascent leading and lagging telomeric strands by labeling them with green and red (respectively) strand-specific PNA probes. T-SCE appears as colocalization of the two probes at the same telomere end (Figure 7). We observed a significant increase in T-SCE events in I/I MEFs compared to control M/M MEFs (Figure 7C), indicating increased homologous recombination at the telomeres of the I/I MEFs, which was not observed in the K/K MEFs of the same PD (10). Such elevated levels of telomeric recombination could be the result of replication stress and replication fork collapse, consistent with the increased TIF formation and telomere fragility.

**Figure 7.**
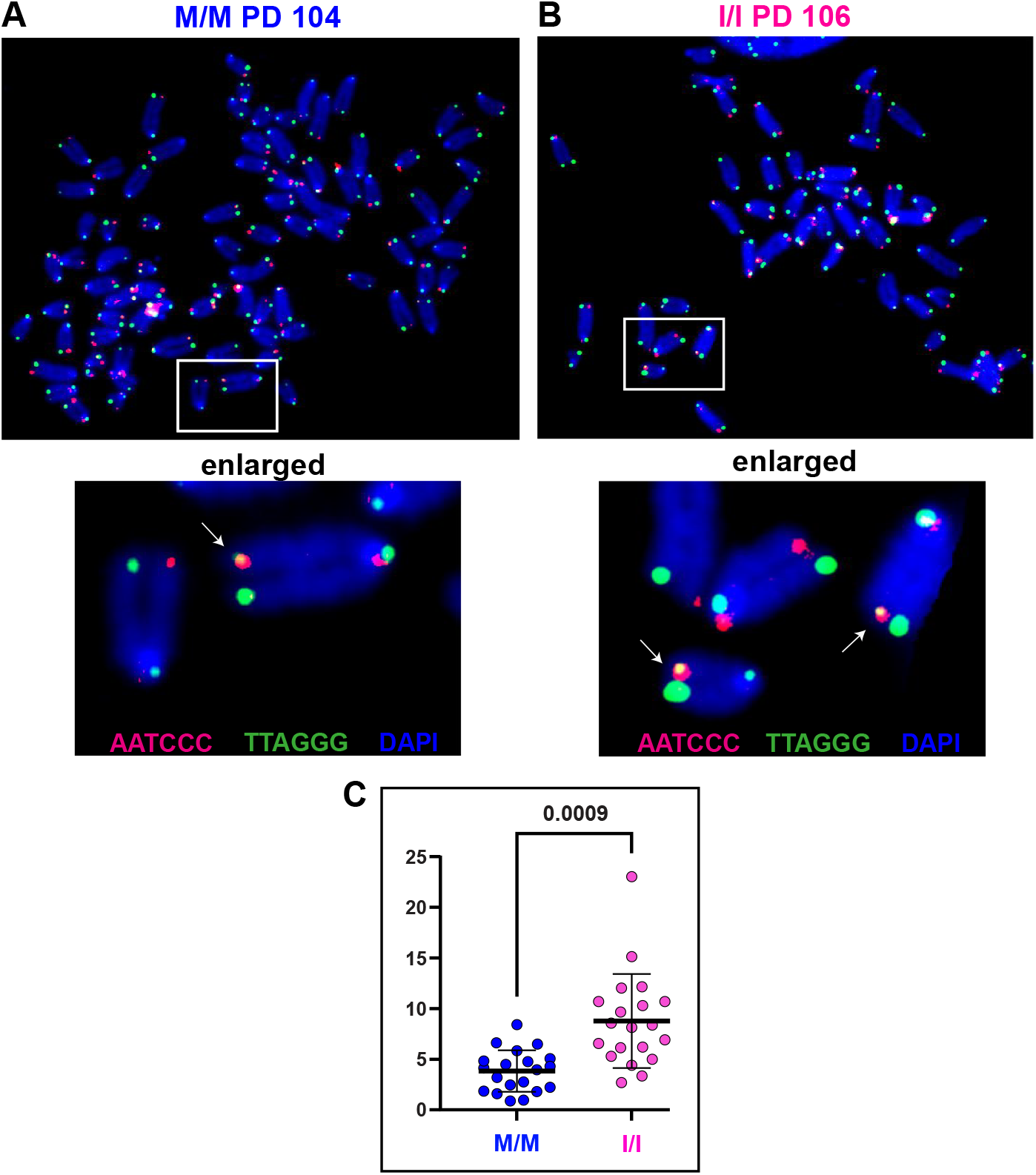
Elevated homologous recombination between sister telomeres in HHS MEFs. (A) Chromatid-Orientation (CO)-FISH was performed on M/M and I/I MEFs at PD106 and PD104, respectively, labeling telomeres with nascent leading and lagging strands with green and red (respectively) strand-specific PNA probes. 20 metaphases were quantified for each culture. Enlarged show examples of Telomere Sister Chromatid Exchange (T-SCE) events. (B) T-SCE events (indicated by arrows) were countified and presented as the percentage from the number of detectable telomeres in each metaphase. Horizontal lines show mean and SD. P-value was calculated by two-tailed paired t-test.

The excision of telomere (t)-loops in the form of telomere (t)-circles has been suggested to cause telomere damage and loss in RTEL1-deficient mouse cells (16,17,39). However, significantly-elevated levels of t-circles were not detected in patient cells harboring the *Rtel1*^M492I^ mutation (9,27). To determine whether increased t-loop excision contributed to the telomere shortening and aberrations observed in the HHS mouse cells, we examined the abundance of t-circle in tail samples by two-dimensional (2D) PFGE (Figure S6). No significant increase in t-circle formation was detected in the HHS mouse cells, as compared to Telomice at the same generation (F5) or to WT control mice, consistent with the observation in patient cells, excluding t-loop excision as a major cause for telomere shortening or other types of telomere damage.

### Why do the two mutations at the same position of RTEL1 have different effects on its function?

The human and mouse RTEL1 proteins display sequence identity higher than 75%. Indeed, using the structure prediction model *Alphafold* (35) we found that the helicase domains of human and mouse RTEL1 are almost identical (Figure 8). The substitution of a methionine by an isoleucine is one of the most conservative substitutions of a protein residue. However, the sulfur atom of a methionine allows for hydrophobic contacts over a longer distance than a carbon atom (5.5Å *versus* 4.5Å) (40). As a result, the methionine interactions with the surrounding residues provide an additional 1-1.5Kcal/mol stabilization to the protein (40). Methionine 492 is buried at the center of the RTEL1 helicase domain, and it is important for its proper folding (Figure 8A). The structural model of the complex of the single-stranded DNA bound to the helicase domain shows two hydrogen bonds between each of the residues K257 and E261 and the DNA substrate (Figure 8B-D). The helix formed by these residues is adjacent to the helix of M492 and makes direct hydrophobic contact with it via V255 and the aliphatic portion of the sidechain of E256 (Figure 8C, D). Replacing M492 with an isoleucine is predicted to abolish the long-range interactions and destabilize this region, affecting DNA binding and processing, and leading a to partially defective RTEL1 helicase core. In addition, the M492I mutation was reported to compromise the interaction of RTEL1 with DNA Polymerase Delta Interacting Protein 3 (PolDIP3) (41). The RTEL1 structural model shows that the edge of the helix that includes M492 is solvent exposed on the opposite side of M492 and therefore accesible for the putative interaction with PolDIP3 (Figure 8E, F). The M492I mutation destabilizes this region, which in turn may subtly affect POLDIP3 binding. This is further supported by our model which places all three residues reported as important for the RTEL1-POLDIP3 interaction (M492, L710 and G739) on the same surface of the RTEL1 protein (41) (Figure 8E, F). A striking difference between the helices formed by M492 in the human and mouse RTEL1 is the presence of a valine residue at position 485 of the human RTEL1 and a leucine in the mouse. An overlay of the human and mouse RTEL1 helicase domains (Figure 8B) shows that the bulkier leucine 485 sidechain displaces the mouse M492 helix further away by 1 Å from the helix containing the DNA interacting residues K257 and E261. The effect of this increased distance on the RTEL1 function is yet to be explored.

**Figure 8.**
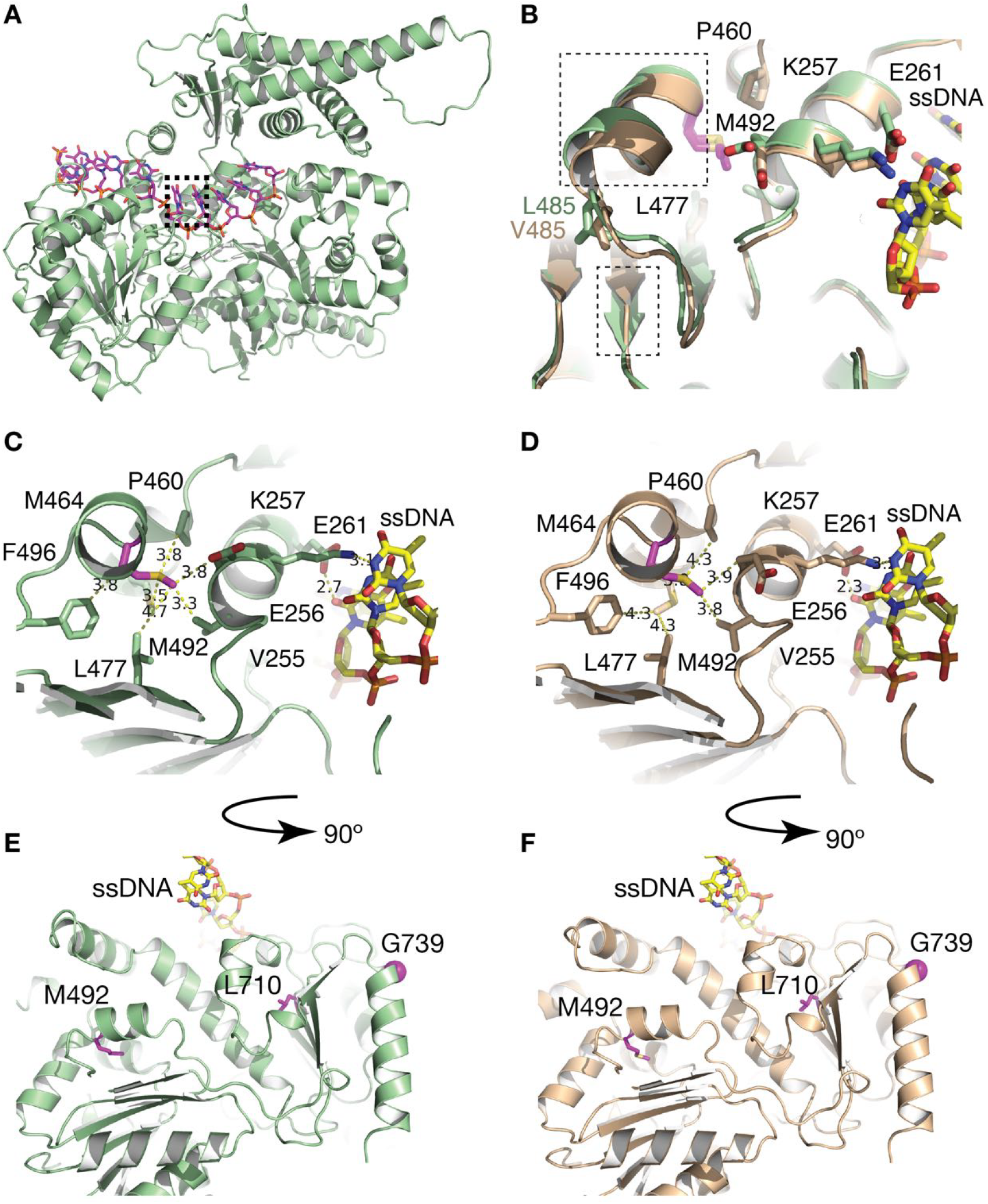
Structural models of the human and mouse RTEL1 helicase domains. (A) A model of the mouse helicase domain of RTEL1 generated by *AlphaFold* (35). (B) Overlay of the region of the human (wheat) and mouse (pale green) RTEL1 helicase domains containing residue 492. Residues V485 and L455, implicated in the subtle reorganization of that region of the protein in the presence of the M492I mutation, and residues K257 and E261, affecting DNA binding, are shown in ‘stick’ format. The displacement of this helix and the adjacent strand between the two species are highlighted in dashed boxes. Interactions of the mouse (C) and human (D) M492 with the surrounding residues, and of K257 and E261 with the single-stranded DNA (ssDNA). The mouse (E) and human (F) RTEL1 structures of (C and D) are rotated 90° to show the positions of M492, L710 and G739, all of which are presumably involved in POLDIP3 binding (41).

## DISCUSSION

RTEL1 has been implicated in diverse pathways essential for genome maintenance and stability, including telomere protection and length regulation (reviewed in (13)). Mutations in *RTEL1* cause diverse symptoms in telomere biology diseases (TBDs) and pulmonary fibrosis, and in cancer predisposition (13). RTEL1-dysfunction in patient cells was associated with severe telomere shortening but also with increased telomeric damage, telomere aberrations and genomic instability, even in cells with normal telomere length (9,23,31). These diverse symptoms and molecular phenotypes associated with RTEL1 deficiency raise the question whether defects other than telomere shortening contribute to disease etiology (13). We have shown previously that changing methionine 492 of *Rtel1* to a lysine in a *M. musculus* mouse model termed ‘Telomouse’ shortened its telomeres to the length of human telomeres (10). The Telomouse telomeres did not display elevated DDR, fragility or other telomere aberrations, except for telomere loss and Robertsonian fusions (10). These observations suggest that the telomouse telomeres are largely protected, unless they shorten below the critical length and induce telomere fusions between the two short (p) arms of the acrocentric mouse chromosomes (Robertsonian fusions), which are tolerated because of the proximity of the two centromeres in the fused dicentric chromosome (42). Thus, the *Rtel1*^M492K^ mutation mostly affected the telomere length regulation function of RTEL1. Here we report on another mouse model, carrying the HHS causing mutation *Rtel1*^M492I^, termed HHS mouse, which exhibits a distinctly different phenotype. The telomeres of HHS mice and MEFs shortened gradually, but to a lesser extent than those of Telomice, as observed by in-gel hybridization and nanopore sequencing (Figures 2, 3 and S5, and Supplementary Tables S1, S2, and S3). Nevertheless, telomere protection in HHS (I/I) MEFs, but not Telomouse (K/K) MEFs, was significantly compromised, as indicated by the increased levels of DDR, telomere fragility, anaphase bridges, micronuclei, and telomere sister chromatid exchange (T-SCE) (Figures 4-7). On the other hand, telomere loss and Robertsonian fusions were elevated in the Telomouse MEFs due to the more severe telomere shortening caused by the M492K mutation (10). Telomere shortening in both Telomouse and the HHS mouse was not associated with increased formation of t-circles (Figure S6), ruling out t-loop excision as a major cause for telomere shortening and aberrations, consistent with the results obtained for HHS patient cells (9,27).

In sum, our results show that the *Rtel1*^M492I^ mutation causes milder telomere shortening but a more severe telomere protection defect than the *Rtel1*^M492K^ mutation, distinguishing two critical functions of RTEL1. We can speculate that the two substitutions have different effects on the structure of the helicase core, the single-stranded DNA binding, and interactions with Poldip3 (Figure 8). Yet, the mechanistic details of how these mutations affect telomere length and telomere protection are yet to be explored. The observations in cells of HHS patients and the HHS mouse support the notion that not only telomere shortening but also length-independent telomere dysfunction contribute to the HHS etiology. While telomere shortening is gradual and may take years to reach a critical short length and cause disease, defects in telomere structure and function may affect physiology more immediately and increase the risk for cancer and pulmonary fibrosis even without significant telomere shortening. Such length-independent effects on telomere function can also explain the more severe symptoms and earlier onset of HHS as compared with dyskeratosis congenita, which is thought to result from telomerase deficiency and telomere shortening. The HHS mouse and Telomouse will serve as invaluable models to study the mechanistic functions of RTEL1 in telomere length regulation and protection, and the consequences of the disruption of these functions on the organismal physiology in TBD, normal aging, and cancer.

## DATA AVAILABILITY

Representative TRF gels are shown in Figures 2 and 3. Additional gels are shown in Supplementary Figure S5. All measured and calculated values are provided in the Supplementary Tables S1 (MEFs) and S3 (mice). The Nanopore telomeric read characteristics are summarized in Supplementary Table S2. The nanopore sequencing data and Telomere Analyzer output folders are available on the OSF repository. Additional data are available upon request to the corresponding authors.

## CODE AVAILABILITY

The code used for filtering and processing Nanopore sequencing reads, termed Telomere Analayzer, is available (34).

## SUPPLEMENTARY DATA

Supplementary Data are available at NAR Online.

## ACKNOWLEDGEMENTS

We are grateful to all members of the Tzfati and Kaestner laboratories for stimulating discussions and assistance, and the members of the Genetically Modified Mouse Core of the Center for Molecular Studies in Digestive and Liver Diseases (P30DK050306) for their help with the derivation of the HHS mouse. We thank Ittai Ben-Porath and Amir Eden for the gifts of WT mice and MEFs; Mark Tigue for the maintenance of the mouse colony; Devora Olam for assistance in mouse blood collection; Adan Mhameed for assistance in quantifying FISH aberrations; Naomi Melamed-Book, Bio-Imaging Unit at The Hebrew University; Dana Avrahami-Tzfati for assistance with CRISPR editing in zygotes; and Ran Avrahami for assistance with statistical analyses.

## FUNDING

This research was supported in parts by the Israel Science Foundation grants 2071/18 and 1342/23 to Y.T., the NIDDK through 1R01DK139049-01 and R01-CA249929 to K.H.K., and by the Israel-UK-Palestine GROWTH Fellowship to R.S.

## CONFLICT OF INTEREST

The authors declare no conflict of interest.

## Notes

### Competing Interest Statement

The authors have declared no competing interest.

### Summary of Updates

The main update to this revised version is the addition of nanopore sequencing of telomeres from the mice and human samples.

