## Supplementary Figures for "Separation of telomere protection from length regulation by two different point mutations at amino acid 492 of RTEL1"

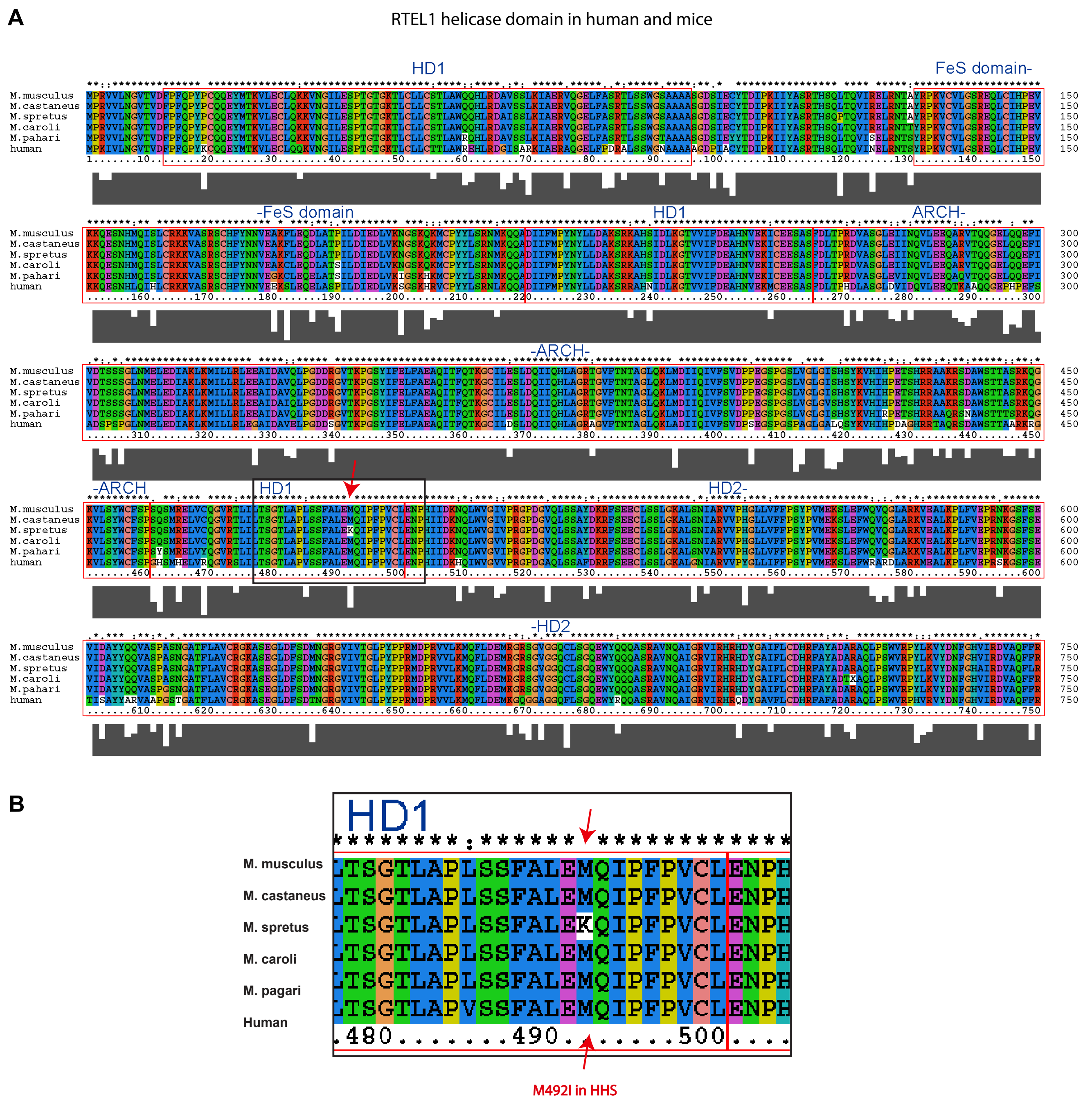


**Figure S1. Conservation of the RTEL1 helicase domain between human and mouse**. (A) Protein sequence alignment of the RTEL1 helicase domain, including HD1, HD2, the iron-sulfur (FeS) and the ARCH domains, showing high conservation between human and different mouse species (*M. musculus, M. castaneus, M. spretus, M. caroli*, and *M. pahari*). (B) The HHS mutation M492I (red arrow) and flanking sequences.

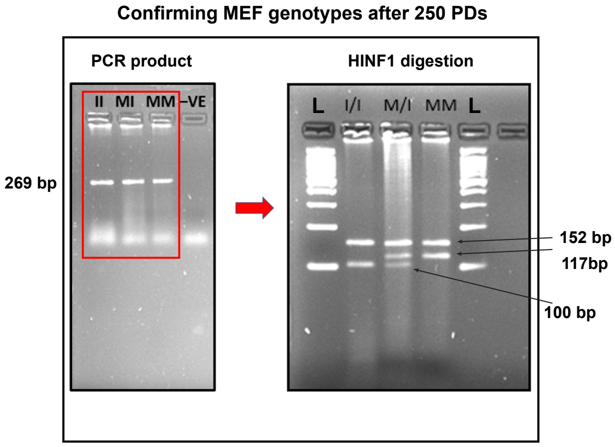


**Figure S2. Genotyping of M/M, M/I, and I/I MEFs.** The genotypes of the MEFs cultures were confirmed at PD 250 by restriction fragment length PCR (RFLP). PCR products were digested with *HinfI* restriction endonuclease. An additional restriction site formed by the M492I mutation shortens the 117 bp fragment by 17 bp.

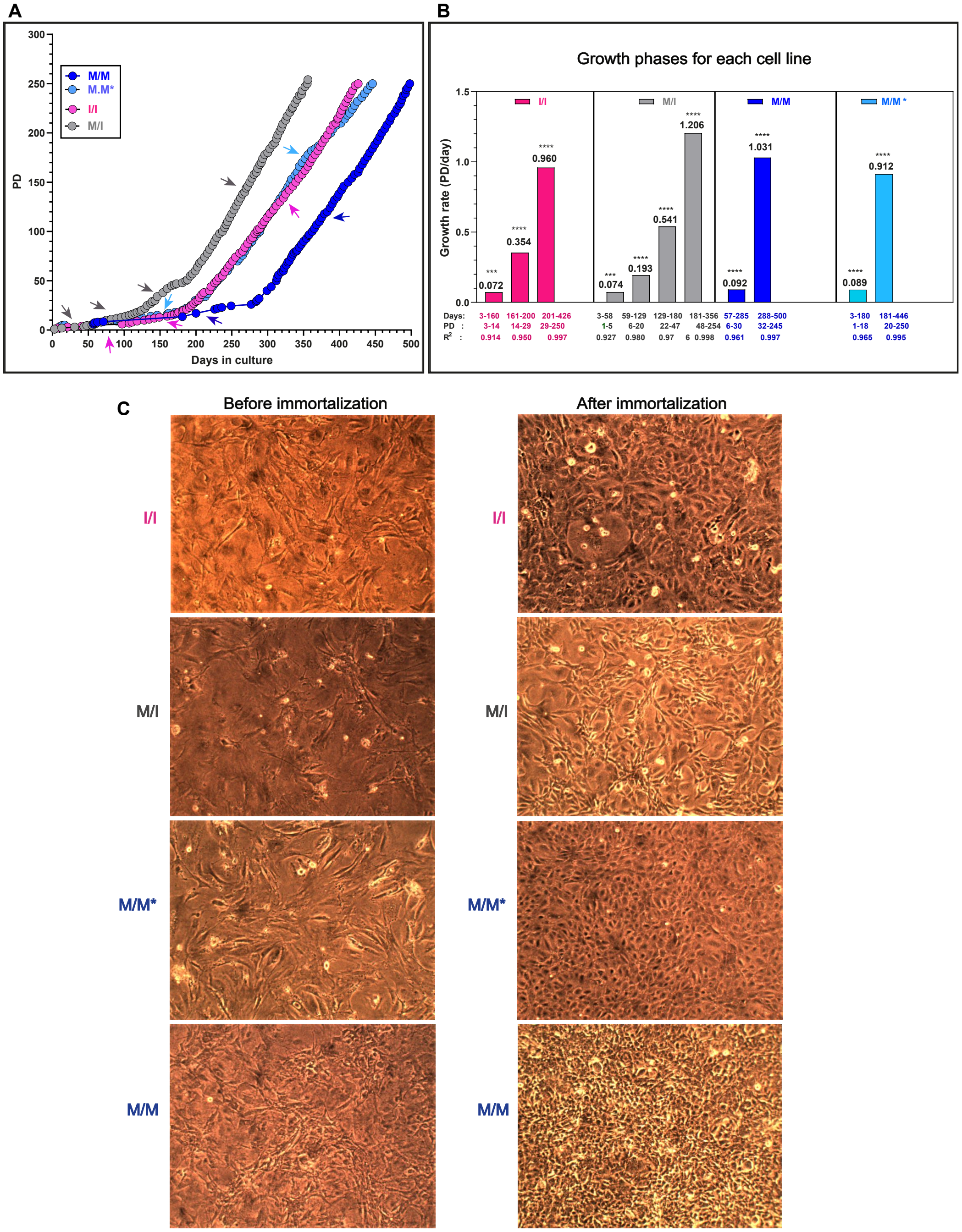


**Figure S3. Immortalization and growth characteristics of MEFs from WT and *Rtel1*$^{M492I}$ mutant mice.** (A) MEFs cultures were prepared from F3 littermate embryos, I/I, M/I and M/M*, generated by mating F2 heterozygous mice carrying the *Rtel1*$^{M492I}$ mutation. Another control M/M MEFs culture (M/M) was generated by intercrossing WT mice which did not have any *Rtel1* mutation in their pedigree, to avoid trans-generational inheritance of short telomeres from the heterozygous mice to the WT offspring. All four cell lines were immortalized by serial passaging and grown to PD 250. Arrows indicate different growth phases which are shown as bars in (B). (B) Growth rates were calculated based on the graphs in (A) and indicated above the bars. Days, PD, and coefficient of determination (R2), for the linear regression line calculated for each growth phase are indicated below. **** indicating P < 0.0001 and *** indicating P<0.001. (C) Representative images show different cell morphology for the different cultures before and after immortalization.

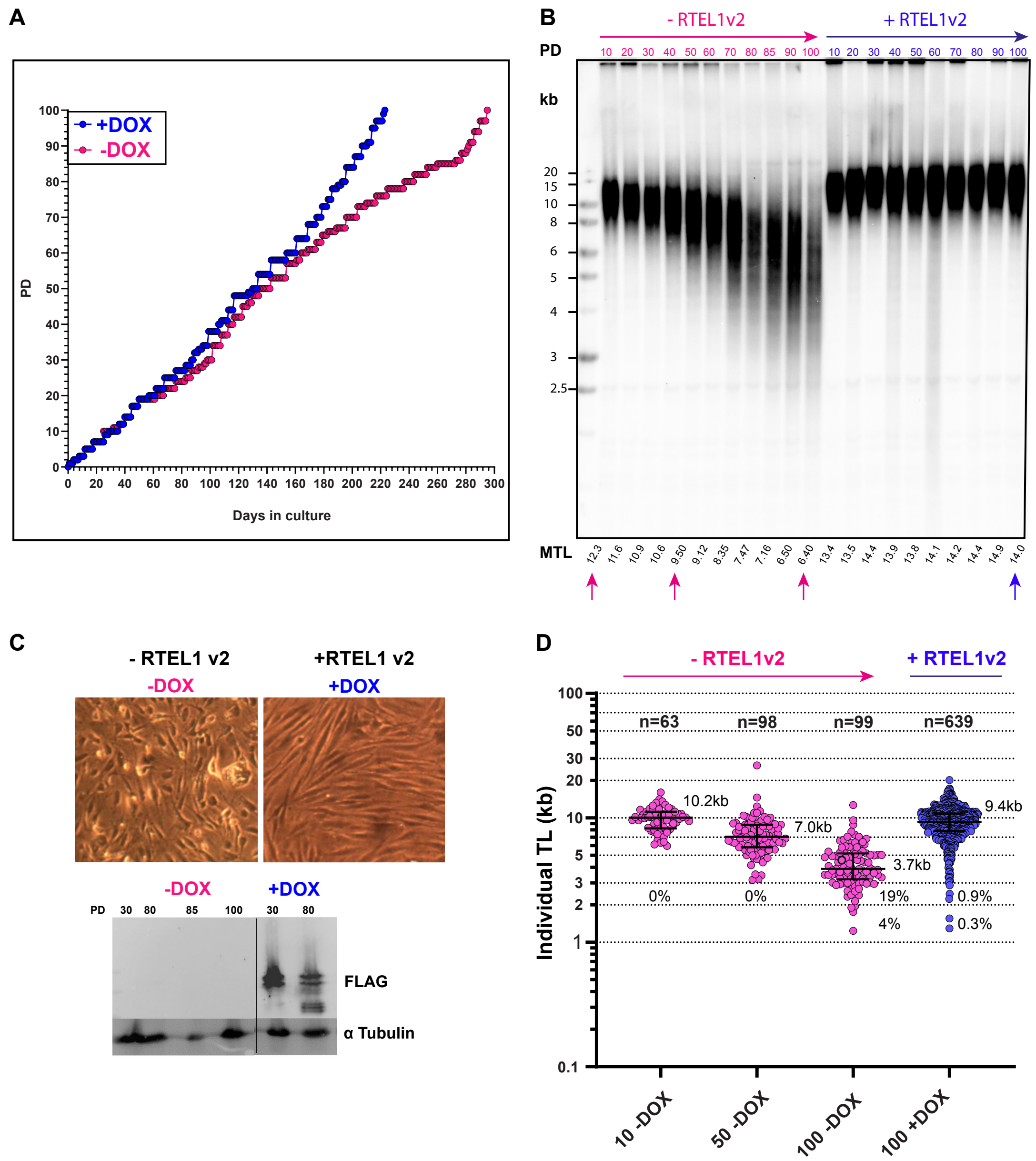


**Figure S4. Silencing the ectopic expression of WT hRTEL1v2 progressively shortened the telomeres of human patient fibroblasts carrying the *RTEL1*$^{M492I}$ mutation.** (A) S2-hTERT-TPP1 fibroblast culture (1) was grown for 58 PD in the presence of 30 ng/ml doxycycline (DOX) to induce the expression of FLAG-tagged RTEL1v2, and then without Dox for 10 PDs. Then the culture was split; half continued to grow in the absence of DOX while the other half was grown with DOX. From this point, PD level was plotted over time in culture till reaching PD 100 with or without DOX. (B) genomic DNA prepared from the S2-hTERT-TPP1-RTEL1v2 cultures (+/-DOX) at the indicated PD were analyzed by in-gel hybridization. Shown is the hybridization of the telomeric probe to the denatured DNA. MTL values, measured by *Telotool* (2), are indicated below the lanes for each sample. (C) Cell morphology and western analysis using anti-FLAG antibody to detect the FLAG-tagged RTEL1v2 in these S2-hTERT-TPP1 fibroblasts. Alpha -tubulin was used as a loading control. (D) The same genomic DNA samples in (B) at the indicated PD were analyzed by *NanoTelSeq*. Shown in scatter plots are lengths of individual telomeres from the +DOX (blue) and -DOX (pink) fibroblast cultures at the indicated PD. Median and quartiles indicated by horizontal lines, and median values on the right and number of telomeric reads above the plots.

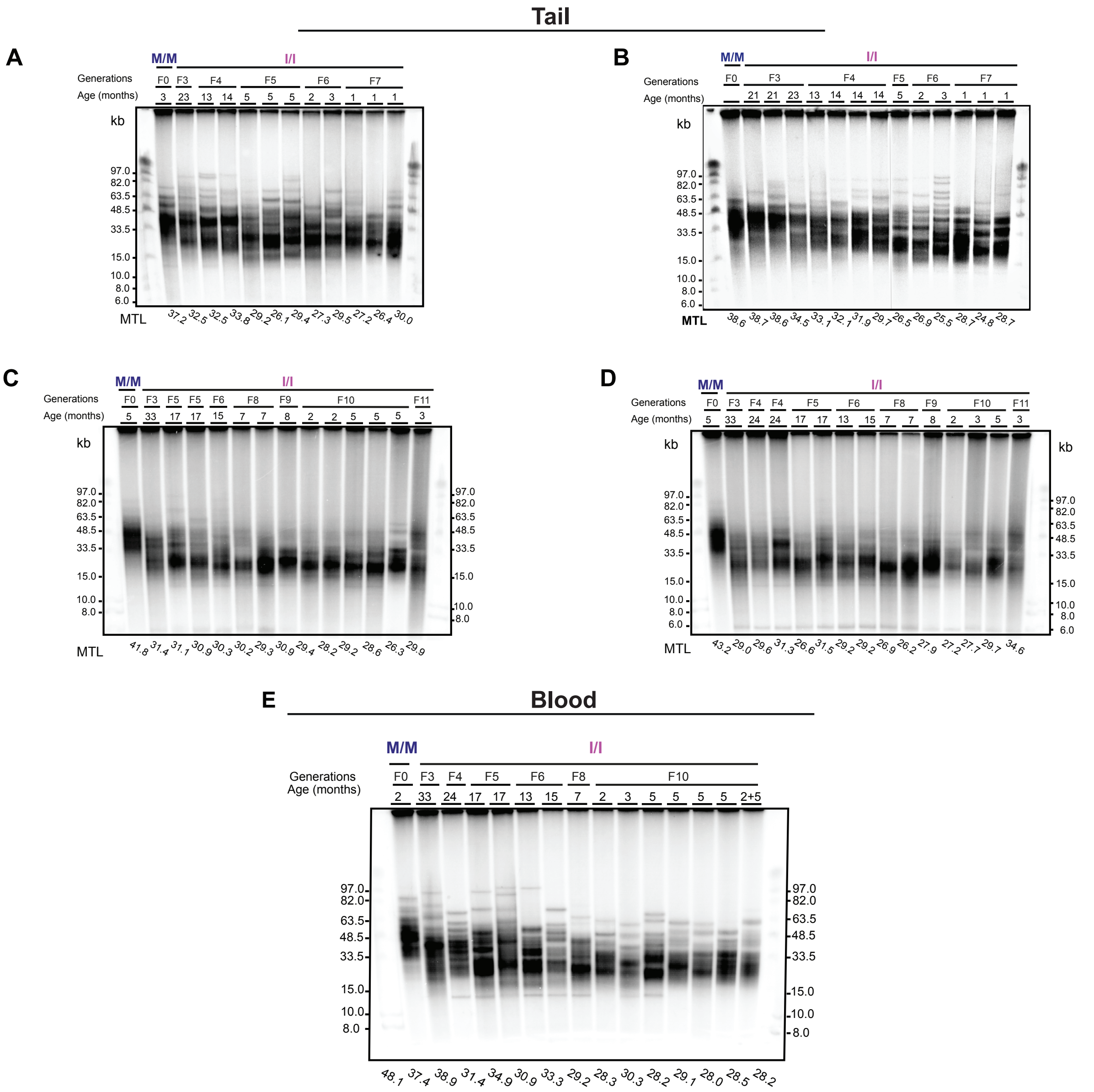


**Figure S5. The HHS mouse telomeres progressively shorten over generations.** Genomic DNA samples extracted from tail (A) or blood leukocytes (B) from mutant (I/I) or WT (M/M) mice at the indicated generations and ages were analyzed by PFGE and in-gel hybridization to the denatured DNA.

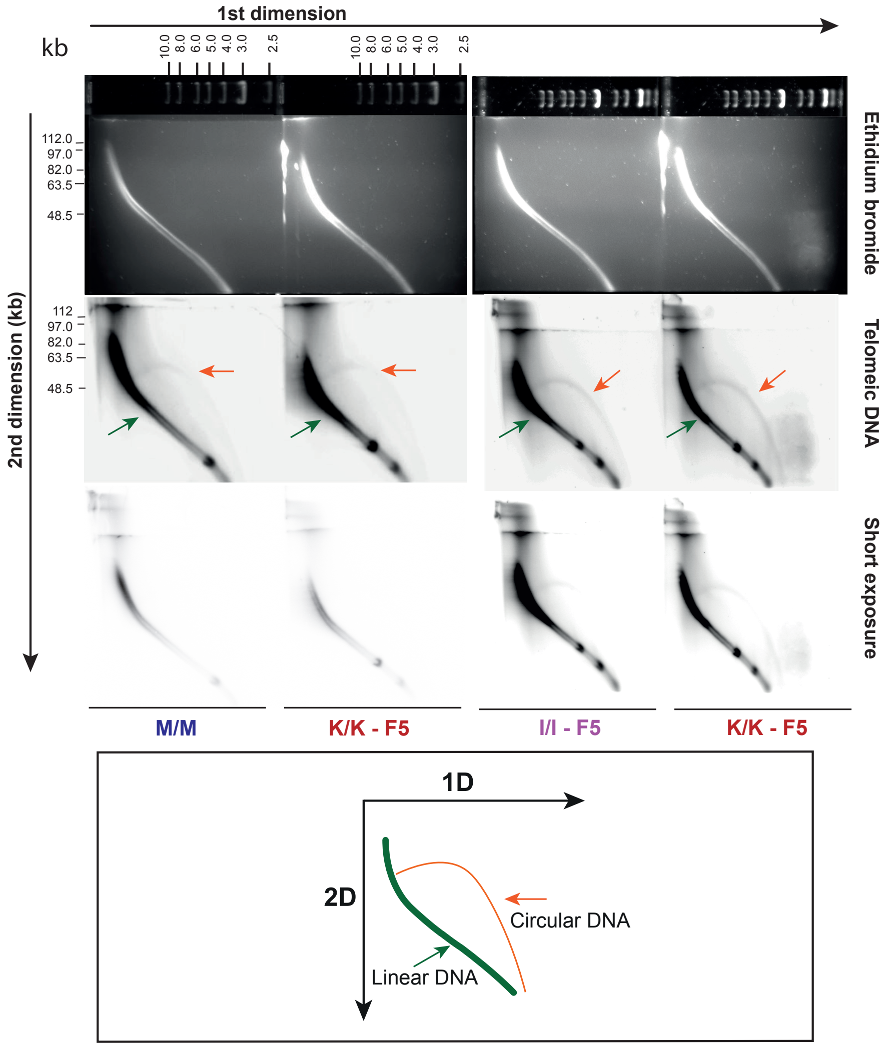


**Figure S6. No increase in t- circle levels in Telomouse and HHS mouse.** (A,B) gDNA (~20-25 µg) extracted from HHS mouse (I/I) and Telomouse (K/K) tail at the same generation (F5), as well as from WT mouse (M/M) was digested with *HinfI* and separated by two-dimensional (2D) PFGE as detailed under Materials and Methods. Shown are two gels, each containing two tail DNA samples: M/M & K/K in (A), and I/I & K/K in (B). The same Ethidium staining shows all genomic DNA fragments. Telomeric circles were detected by in-gel hybridization with a C-rich radioactive probe and visualized using a Phosphor-Imager. Short exposure shows comparable amounts of telomeric DNA in each sample. The green and red arrows indicate linear- and circular-form of telomeric DNA, respectively. (C) A scheme showing the 2D gel principle.

### SUPPLEMENTARY FIGURES REFERENCES

1. Awad, A., Glousker, G., Lamm, N., Tawil, S., Hourvitz, N., Smoom, R., Revy, P. and Tzfati, Y. (2020) Full length RTEL1 is required for the elongation of the single-stranded telomeric overhang by telomerase. *Nucleic Acids Res*, **48**, 7239-7251.

2. Gohring, J., Fulcher, N., Jacak, J. and Riha, K. (2014) TeloTool: a new tool for telomere length measurement from terminal restriction fragment analysis with improved probe intensity correction. *Nucleic Acids Res*, **42**, e21.
