## Supplementary Tables Legends for "Separation of telomere protection from length regulation by two different point mutations at amino acid 492 of RTEL1"

**Supplementary Table 1. Mouse embryonic fibroblasts.** Genomic DNA samples, prepared from WT (M/M, M/M*), homozygous mutant (I/I) or heterozygous (M/I) for the *Rtel1*^M492I^ mutation, were grown over 250 population doublings (PD), sampled at different PD, and analyzed in ten different gels by PFGE and in-gel hybridization to the denatured DNA. Mean telomere length was quantified using *TeloTool* {Gohring, 2014 #3425} using the ‘corrected’ mode. Percentage of telomeres below 15 kb (TL< 15) or 20 kb (TL<20) was calculated from the mean TRF length and the standard deviation (SD).

**Supplementary Table 2. Nanopore sequencing for MEFs and mice**

Undigested high molecular weight genomic DNA samples prepared from the Rtel1 I/I MEFs at PD250, K/K MEFs at PD250 , M/M MEFs at PD250, WT mouse blood 13 months old, Telomouse F14 blood and tail 13 months old, and HHS mouse F14 blood and tail 13 months old were sequenced by NanoTelSeq. Telomeric reads were identified and filtered as detailed in the methods.

**Supplementary Table 3. Mice.** Genomic DNA samples from 48 I/I and 24 M/M (100% pure Blk6 C57BL/6) mice were analyzed multiple times in separate gels PFGE (as indicated) and in-gel hybridization to the denatured DNA. Mean telomere length was quantified using *TeloTool* {Gohring, 2014 #3425} using the ‘corrected’ mode. Percentage of telomeres below 15 kb (TL< 15) or 20 kb (TL<20) was calculated from the mean TRF length and the standard deviation (SD). The average age for I/I mice analyzed was 329.0 days, while the average age for M/M mice was 355.5 days. At each specific generation, the following mice were analyzed:

F0 (M/M): 24 mice (63 - 665) days old

F2 (I/I): two mice 355 and 433 days old

F3 (I/I): 6 mice (115 – 1014) days old

F4 (I/I): 7 mice (174 – 738) days old

F5 (I/I): 5 mice (149 – 505) days old

F6 (I/I): 4 mice (48 – 455) days old

F7(I/I): 6 mice 38 days old

F8 (I/I): 5 mice 215 days old

F9 (I/I): 2 mice 239 days old

F10 (I/I):10 mice (53 – 154) days old

F11 (I/I): one mouse 80 days old
